# Root membrane ubiquitinome under short-term osmotic stress

**DOI:** 10.1101/2021.12.06.471398

**Authors:** Nathalie Berger, Vincent Demolombe, Sonia Hem, Valérie Rofidal, Laura Steinmann, Gabriel Krouk, Amandine Crabos, Philippe Nacry, Lionel Verdoucq, Véronique Santoni

## Abstract

Osmotic stress can be detrimental to plants, whose survival relies heavily on proteomic plasticity. Protein ubiquitination is a central post-translational modification in osmotic mediated stress. Plants use the ubiquitin (Ub) proteasome system to modulate protein content, and a role for Ub in mediating endocytosis and trafficking plant plasma membrane proteins has recently emerged. In this study, we used the K-ε-GG antibody enrichment method integrated with high-resolution mass spectrometry to compile a list of 719 ubiquitinated lysine (K-Ub) residues from 450 *Arabidopsis* root membrane proteins (58% of which are transmembrane proteins), thereby adding to the database of ubiquitinated substrates in plants. Although no Ub motifs could be identified, the presence of acidic residues close to K-Ub was revealed. Our ubiquitinome analysis pointed to a broad role of ubiquitination in the internalization and sorting of cargo proteins. Moreover, the simultaneous proteome and ubiquitinome quantification showed that ubiquitination is mostly not involved in membrane protein degradation in response to short osmotic treatment, but putatively in protein internalization as described for the aquaporin PIP2;1. Our in silico analysis of ubiquitinated proteins shows that two E2 Ub ligases, UBC32 and UBC34, putatively target membrane proteins under osmotic stress. Finally, we revealed a positive role for UBC32 and UBC34 in primary root growth under osmotic stress.

## Introduction

Plants are exposed to different types of abiotic stress conditions such as drought or salinity that result in diminished plant growth and crop productivity (Lobell et al., 2011). Most of these conditions impose osmotic stress on plants by reducing the water potential of the environment. The consequences of osmotic stress manifest as inhibited cell elongation, stomata closure, reduced photosynthetic activity, the translocation of assimilates, changes in various metabolic processes, and disturbances in water and ion uptake.

The ability of plants to survive these abiotic stresses relies heavily on their proteomic plasticity. Protein stability, activity, localization, and interactions with partners have all been widely described as governed by ubiquitination (Mukhopadhyay and Riezman, 2007; Nelson and Millar, 2015). Ubiquitin (Ub) is a 76-amino acid polypeptide that is highly conserved in eukaryotes and is ubiquitously found in tissues. It is linked to either target proteins or itself through the sequential action of three enzyme classes: Ub-activating enzymes (E1s), Ub-conjugating enzymes (E2s), and Ub ligases (E3s) (Callis, 2014). The activities of these enzymes ultimately result in the covalent attachment of Ub to a lysine (K) residue in the target protein. Ubiquitination can result in the conjugation of a single moiety (mono-ubiquitination), multiple Ub molecules that are individually attached (multi-mono-ubiquitination), or in the form of a chain (poly-ubiquitination) attached to a specific substrate. Poly-Ub chains are formed by the further attachment of Ub moieties linked together by one of the seven lysine residues present in a Ub molecule (K6, K11, K27, K29, K33, K48, and K63), or by the N-terminal methionine in the form of head-tail linear repeats (Pickart and Eddins, 2004). Poly-Ub chains exhibit different topologies and are associated with diverse biological functions (Walsh and Sadanandom, 2014). As an example, poly-ubiquitination involving residue K48 from ubiquitin (K48-Ub linkage) triggers the degradation of target proteins by the 26S proteasome (Pickart and Eddins, 2004). Much less is known about the other poly-ubiquitination chain linkages (Pickart and Eddins, 2004; Walsh and Sadanandom, 2014). The K63-Ub linkage has been widely studied in yeast and mammals and more recently in plants, and includes roles in the endocytosis of plasma membrane proteins, DNA damage responses and, to a lesser extent, autophagy and signaling (Komander and Rape, 2012; Romero-Barrios et al., 2020).

A role for ubiquitination in abiotic stress tolerance has emerged from the study of transgenic plants overexpressing Ub genes that become more tolerant to multiple abiotic stresses (Guo et al., 2008; Kang et al., 2016), and from the expression analysis of E2s and E3s. The *Arabidopsis* genome contains over 1,400 genes encoding E3s, 37 canonical E2s, and seven E2 variant proteins (Stone, 2018). The number of E3s associated with abiotic stress has increased dramatically over the last decade (Stone, 2018). In particular, exposure to stress (such as osmotic stress) increases abscisic acid levels; moreover, the number of E3s associated with regulating abscisic acid production, signaling, and response continues to grow, now including at least 25 different E3s (Stone, 2018). In addition to E3s, it appears that E2s are not only utilized as Ub-transferring components, but also play an active role in regulating the ubiquitination pathway (Zhou et al., 2010; Cui et al., 2012; Chen et al., 2016). Furthermore, E2 enzymes can interact directly with their targets (Liu et al., 2012; Pan et al., 2020). Indeed, recent studies have suggested that E2s regulate the specificity of target ubiquitination (Turek et al., 2018; Romero-Barrios et al., 2020).

The low stoichiometry and short lifespans of ubiquitinated proteins present obstacles to the identification of ubiquitinated proteins. K-ε-GG (DiGly, the remnant from ubiquitinated proteins following trypsin digestion) antibody affinity enrichment provides an efficient method to gain proteome-wide insight into ubiquitination processes, by capturing and concentrating this remnant of ubiquitinated proteins treated with trypsin prior to MS/MS (Xie et al., 2015; Guo et al., 2017; Zhang et al., 2017; Wang et al., 2019; He et al., 2020; Grubb et al., 2021). In the present study, we used a high-affinity K-ε-GG antibody enrichment technique combined with high-accuracy MS/MS analyses to generate quantitative ubiquitinome and proteome profiles of root membrane proteins from plants treated with a short-term osmotic treatment. Altogether, we provide an extensive inventory of K-Ub residues in a membrane protein fraction, highlight the presence of acidic residues in the vicinity of the K-Ub residue, reveal a role for ubiquitination outside of the membrane protein degradation process in response to short-term osmotic treatment, and demonstrate that the E2 ubiquitin ligases UBC32 and UBC34 are positive regulators of primary root growth during osmotic stress.

## Results

The ubiquitinome response to osmotic stress was investigated by treating plants with 200 mM mannitol for 1 h, followed by a combined quantitative analysis of the proteome and ubiquitinome of a microsomal fraction, according to the proteomic workflow described in Figure 1.

**Figure 1.**
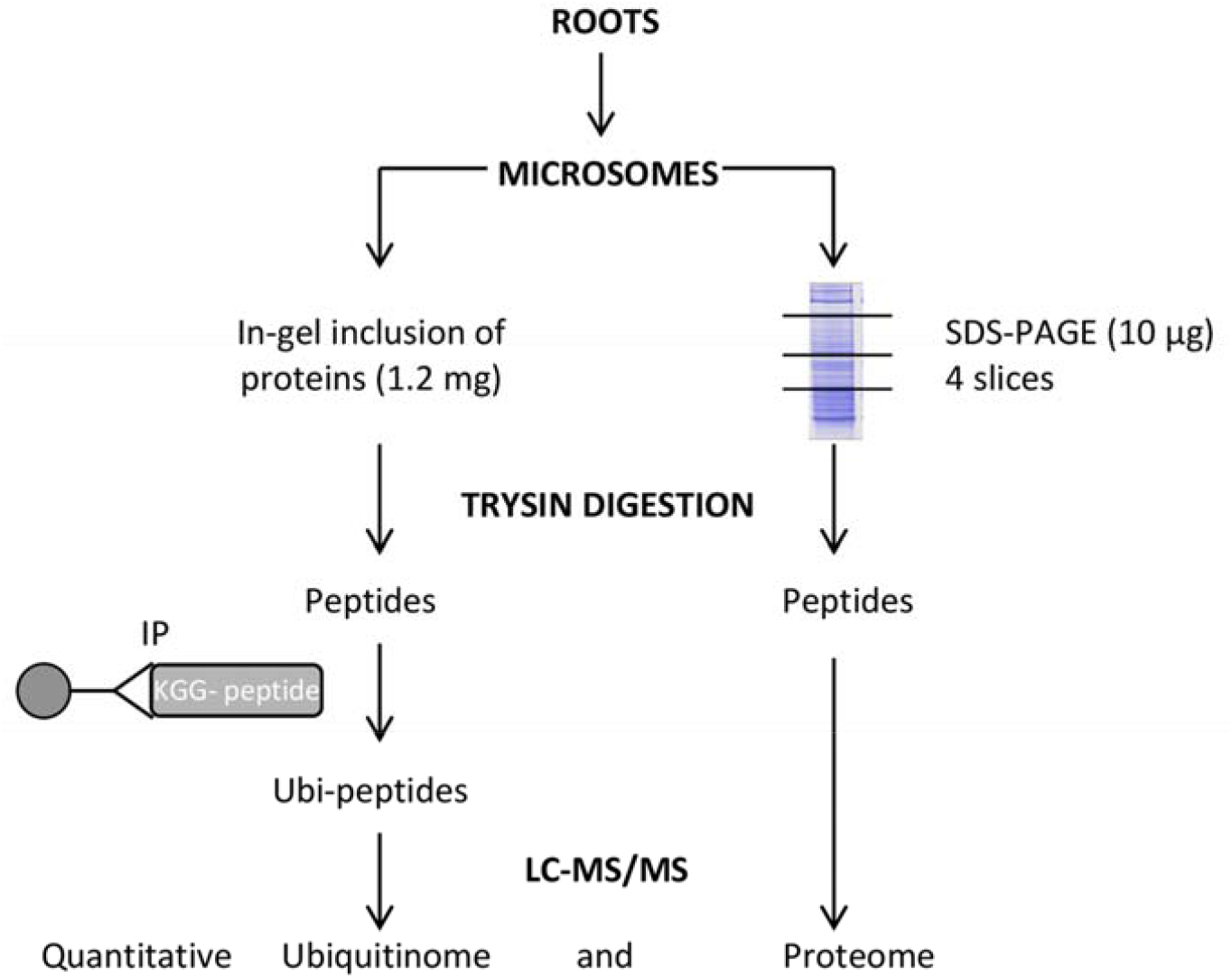
Workflow for quantitative profiling of the proteome and the ubiquitinome in Arabidopsis root membrane proteins upon mannitol treatment. LC-MS/MS: liquid chromatography-tandem mass spectrometry. IP: Immunopurification.

### Differentially accumulated proteins in response to mannitol treatment

A total of 6,081 proteins were identified based on an identification with at least 2 peptides (Table S1), of which 26% were transmembrane proteins (Table S1). Using GO analysis of cell component terms, we showed that a majority of GO terms were associated with membrane proteins, even when the extrinsic proteome (*i*.*e*. proteins without any transmembrane domain) was exclusively considered (Figure S1). These results show that the microsomal fraction is enriched in membrane proteins. Treating plants with 200 mM mannitol for 1 h resulted in differentially accumulated proteins (DAPs), which were identified through a quantitative label-free approach. Among the 226 DAPs that were identified, 132 were up-accumulated (average increase: 1.51x), and 94 were down-accumulated (average decrease: 0.61x) (Table S2). In addition, 1 protein appeared upon mannitol treatment, while 10 proteins disappeared (Table S2). DAPs were classified according to the GO functional categories of “biological process”, “molecular function”, and “cellular component” (Figure S2). Interestingly, enriched functions mostly concerned ATPase activities (Figure S2), in agreement with observations showing the tight regulation of plasma membrane H^+^-ATPase in response to several biotic and abiotic stress responses (Falhof et al., 2016).

### Characterization of the root membrane ubiquitinome

To identify ubiquitinated proteins in *Arabidopsis* roots, we combined immunoaffinity enrichment (using a high quality anti-K-ε-GG antibody; PTM biolabs) and high-resolution mass spectrometry. Ubiquitinated peptides (Ubi-peptides) were considered as long as they had a score > 40 and if they were identified in at least two independent samples. 719 Ubi-peptides harboring a total of 786 K-Ub residues belonging to 450 proteins were identified, 264 of which contained at least one transmembrane domain (Table S3, Table S4). Our GO enrichment analysis showed that ubiquitinated proteins were enriched in transporters, in proteins involved in the regulation of intracellular pH, and in cellular trafficking processes that were characterized by the GO terms “vesicle budding from membrane”, “clathrin-dependent endocytosis” and “membrane invagination” (Figure 2) and included 15 proteins (Table S3). These observations suggest that membrane proteins, as well as the proteins that drive their trafficking are ubiquitinated.

**Figure 2.**
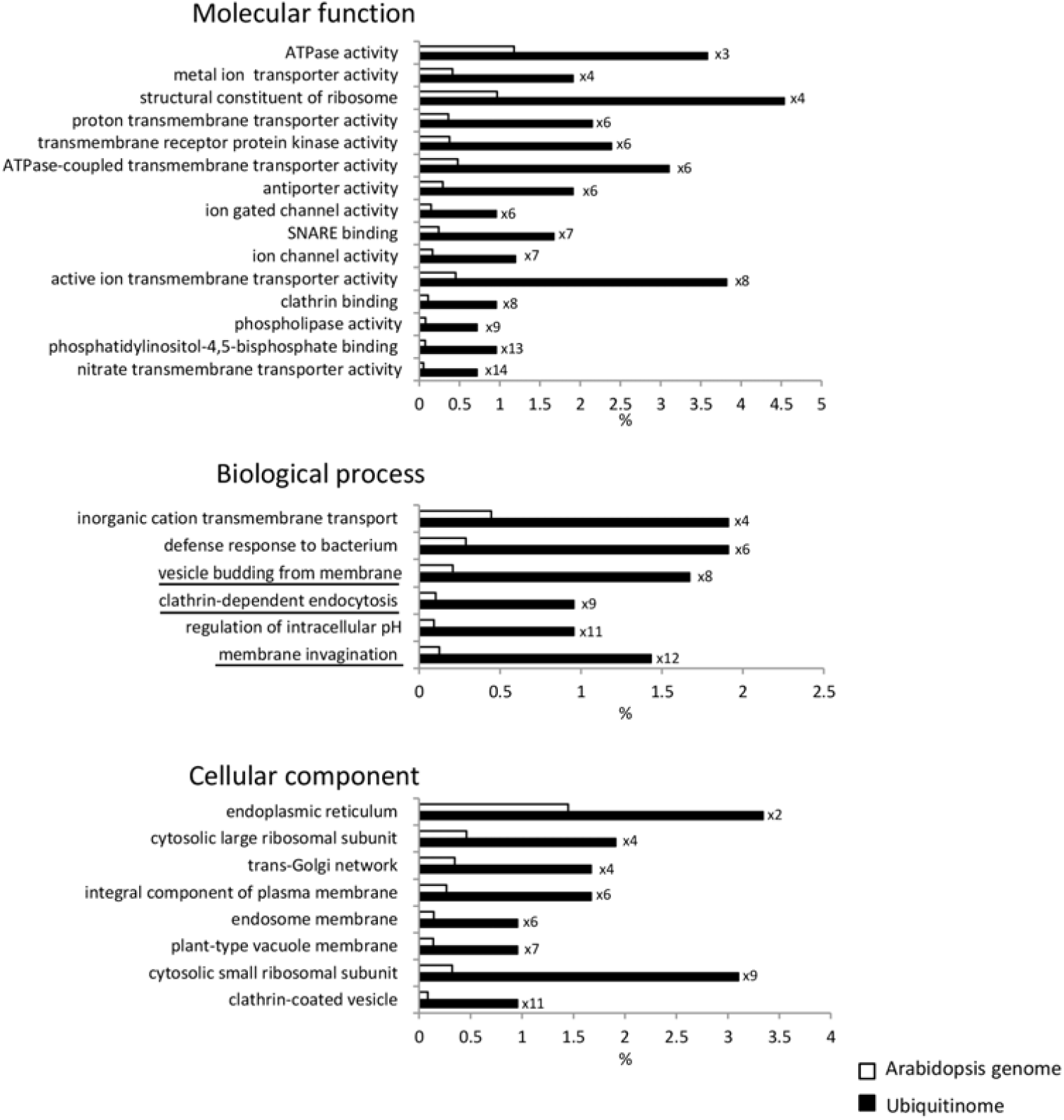
Functional enrichment analysis of ubiquitinated root microsomal proteins. The percentage is calculated with regard to the number of identified ubiquitinated proteins (black) and the total number of Arabidopsis proteins (white). Numbers indicate the fold enrichment by comparison to the Arabidopsis genome. Underlined biological processes concern proteins involved in intracellular trafficking, and include 15 genes (Table S3).

Consensus peptide motifs for K-Ub residues were extracted using p-logo (O’Shea et al., 2013). In total, 643 unique ubiquitinated sites were unable to highlight one unique motif (Table S5, Figure 3A-B). However, the presence of an acidic amino acid close to K-Ub was observed in all motifs except one. Although the nature of the poly-Ub linkage determines the role of ubiquitination, this information is not available in a bottom-up proteomics strategy in which the use of trypsin induces Ub proteolysis from ubiquitinated proteins. However, ubiquitinated peptides arising from the Ub protein itself can provide information about poly-Ub linkages that occur in a protein sample. Seven K-

**Figure 3.**
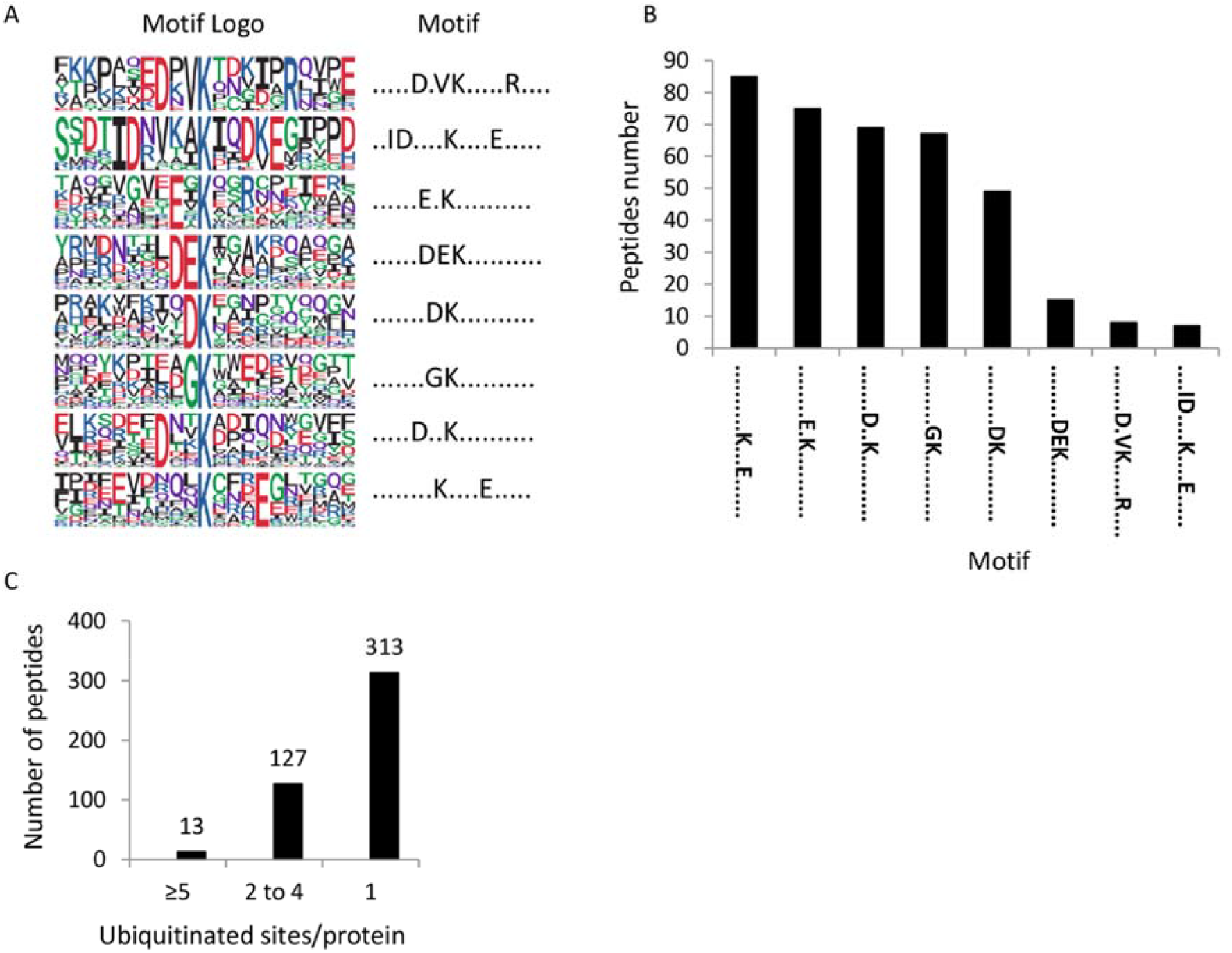
Motif analysis of identified K-Ub residues in root microsomes. A. Ubiquitination motifs and the conservation of K-Ub residues. The height of each letter corresponds to the frequency of the amino acid residue in that position. The central K refers to the K-Ub residue. B. The number of identified peptides containing a K-Ub residue in each motif. C. The number of K-Ub sites per protein.

Ubs were thus identified (K6, K11, K27, K29, K33, K48, K63) within Ub (Figure S3), showing multiple Ub-linkages within membrane proteins. Although peptide intensity is not indicative of the absolute quantity of each Ubi-peptide, the K48- and K63-Ub linkages appeared to predominate the poly-Ub linkages (Figure S3). One way to confirm these observations would be with a targeted proteomics approach that can provide absolute quantification of peptides (Tsuchiya et al., 2013). Nevertheless, our results reveal that mannitol treatment does not significantly modify the proportion of each poly-Ub linkage (Figure S3).

### Differentially accumulated ubiquitinated proteins in response to mannitol treatment

Out of 374 quantified Ubi-peptides, 82 showed quantitative variations in which 54 Ubi-peptides were up-accumulated and 28 were down-accumulated (Table S6). Enrichment-based clustering analyses showed that the ubiquitination of proteins altered by mannitol treatment mainly concerns ATPase, transporters and SNARE binding activities (Figure S4). An inverse quantitative relationship between a protein’s abundance and its ubiquitinated form could be indicative of a role for ubiquitination in protein degradation. However, none of the 43 up-accumulated Ubi-peptides were affiliated with a decreased abundance in the corresponding protein (Table 1). In addition, among 24 down-accumulated Ubi-peptides, only 6 of them corresponded to accumulated proteins including CARK1, HIR2, NRT2;1, PIRL5, AT1G48210.2 and AT3G47210.1 (Table 1). Thus, a role for ubiquitination in degrading these protein could be considered. By contrast, for a majority of the proteins, the absence of any inverse quantitative relationship between the protein and its ubiquitinated form suggests that upon short-term osmotic treatment, ubiquitination could interfere with protein function or cellular localization rather than with protein stability.

**Table 1.**
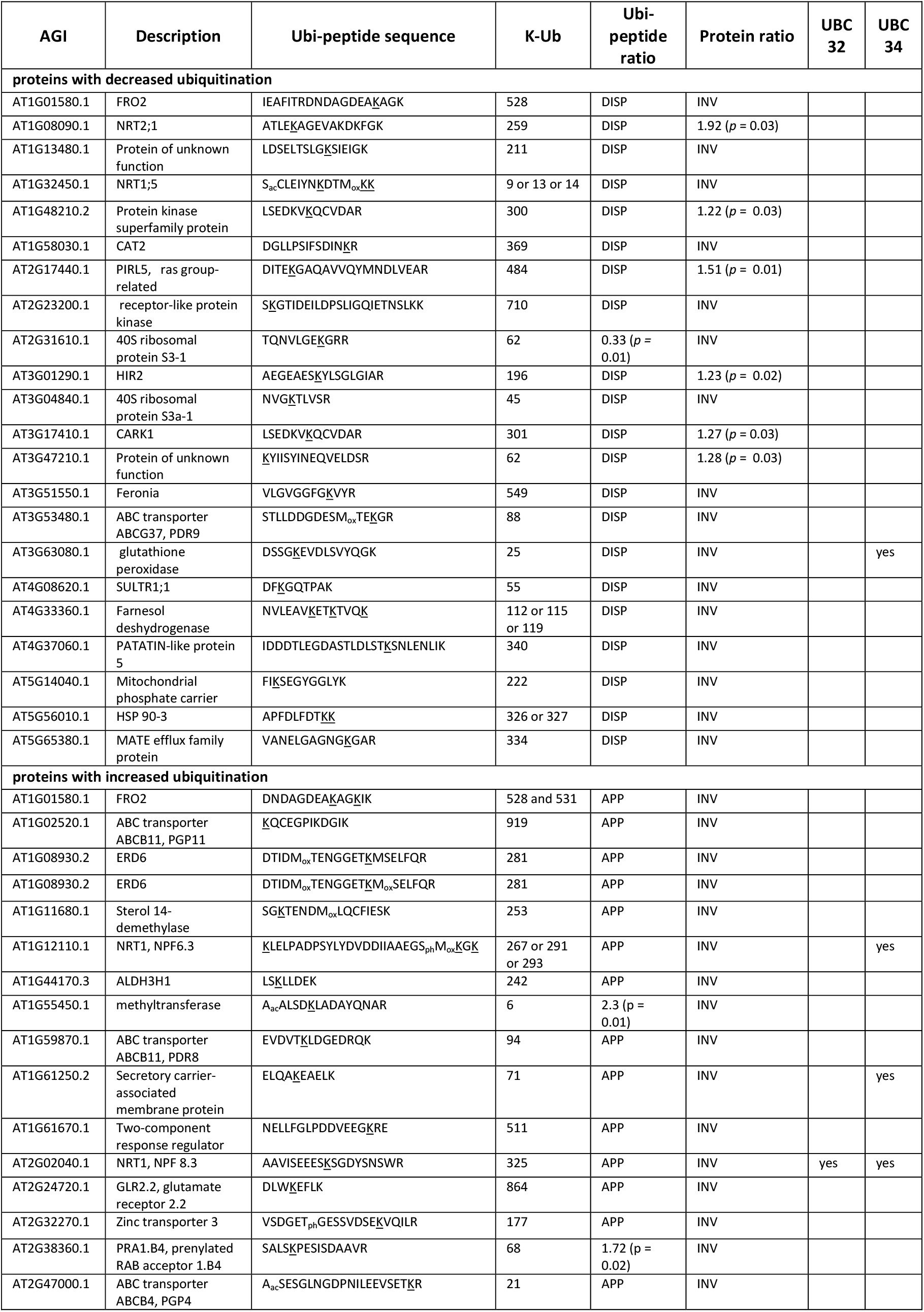
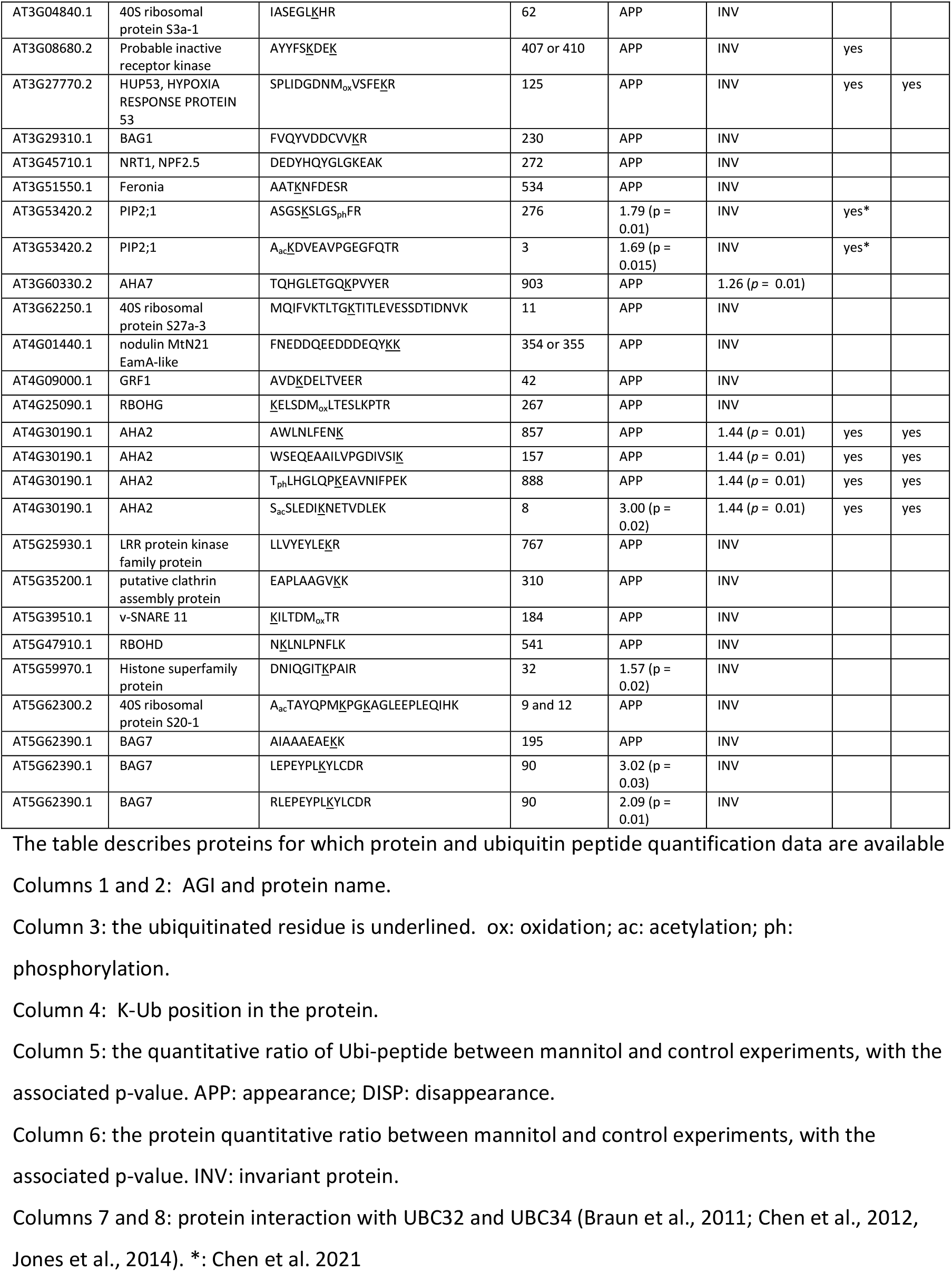
Variations in the ubiquitinated peptide and the corresponding protein in response to mannitol treatment

### Ubiquitination of PIP aquaporins

Aquaporins define a large family of ubiquitous integral membrane proteins that mediate the transport of water and small neutral solutes across membranes (Maurel et al., 2015). Plant aquaporins show a large variety of isoforms. Indeed, 35 homologs within four homology subclasses have been identified in *Arabidopsis*. The plasma membrane intrinsic proteins (PIPs) consist of 13 isoforms further subdivided into the PIP1 and PIP2 subgroups, and are the most abundant aquaporins in the plasma membrane (Johanson and Gustavsson, 2002; Quigley et al., 2002). All 13 members of the PIP family were identified in this study (Table S1), and 9 of them exhibited ubiquitinated residues in their N- and/or C-terminus (Table S3, Figure S5). Since only 3 lysine residues are described in the literature as ubiquitinated (Chen et al., 2021; Grubb et al., 2021), the present work greatly increases our knowledge regarding the ubiquitination of PIPs. Increased ubiquitination of K3 and K276 was observed in PIP2;1 upon short-term mannitol treatment (Table 1). K276 ubiquitination was recently shown to mediate PIP2;1 degradation upon long-term drought (Chen et al., 2021). However, since we observed that PIP2;1 cellular abundance was unchanged (Table 1), the increased ubiquitination of K276 observed when plants are subjected to a 1-h mannitol treatment cannot be involved in PIP2;1 degradation.

Next, we checked if there is a role for K3 ubiquitination using a simplified system of *Arabidopsis* suspension cells with a low basal level of endogenous PIPs. We overexpressed PIP2;1 either wild-type or carrying point mutations at K3 in alanine (K3A) and in arginine (K3R) (Santoni et al., 2006) with the aim of preventing ubiquitination at that site. We previously showed by western blot analysis of total protein extracts using an anti-PIP2;1 peptide antibody that there is a significantly strong overexpression of PIP2;1 in these cells, as compared to untransformed cells or cells transformed with an empty vector (PG) (Santoni et al., 2006). Here, using an ELISA assay, we observed a significant accumulation of PIP2;1 in suspension cells overexpressing PIP2;1-K3A and PIP2;1-K3R as compared to PIP2;1-WT (Figure 4), suggesting that K3 could be a ubiquitinable residue participating in the degradation of PIP2;1. However, in roots placed under short-term osmotic treatment, increased K3 ubiquitination did not correlate with decreased PIP2;1 abundance, suggesting (as for K276) an additional role for K3 ubiquitination outside of PIP2;1 degradation.

**Figure 4.**
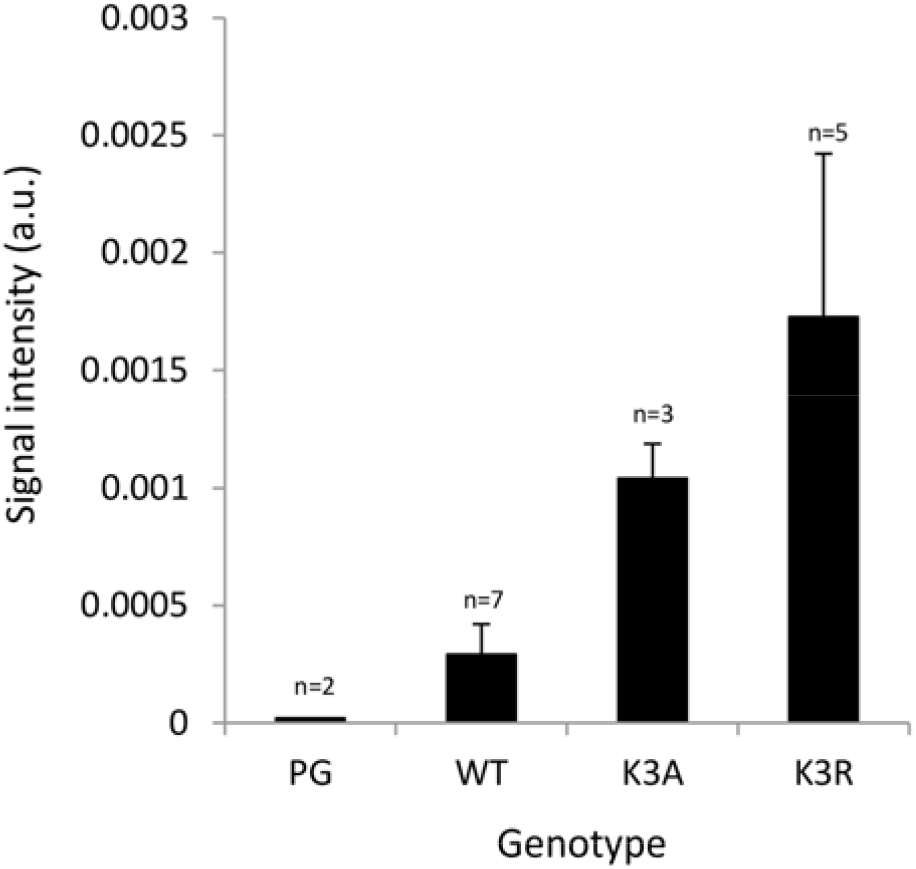
Relative abundance of PIP2 isoforms in Arabidopsis suspension cells overexpressing PIP2;1 WT and carrying a point mutation of K3 in alanine (K3A) and in arginine (K3R). ELISA assays were performed with total protein extracts and anti-PIP2 antibody (Santoni et al., 2006). The number of independent stable cell lines is indicated. Data were from three individual ELISA assays per cell line. Standard error is shown.

### The interactome of ubiquitinated proteins

For additional insight into the extent of the role of ubiquitination, we next constructed a network for the ubiquitinated proteins identified in this work and their interactants identified in previous yeast-two hybrid (Braun et al., 2011) and Split-Ub global approaches (Chen et al., 2012; Jones et al., 2014). This final network consisted of 1,011 proteins (Table S7a, Figure 5). Transport and trafficking functions were enriched in such interactome and, interestingly, the most enriched process concerns ubiquitination, with the GO term “protein K63 linked ubiquitination” showing a 37-fold enrichment (Figure S6). Five E2s (UBC26, UBC32, UBC34, UBC35, UEVD1) and one E3 (UPL6) were identified in this network (Table S7a). We investigated the extent of interaction of these enzymes with the ubiquitinated proteins identified in the present work, and identified two E2s, UBC32 and UBC34, which putatively interact with 15 and 31 ubiquitinated transmembrane proteins involved in water and ion transport, respectively (Figure 5; Table S7b-c).

**Figure 5.**
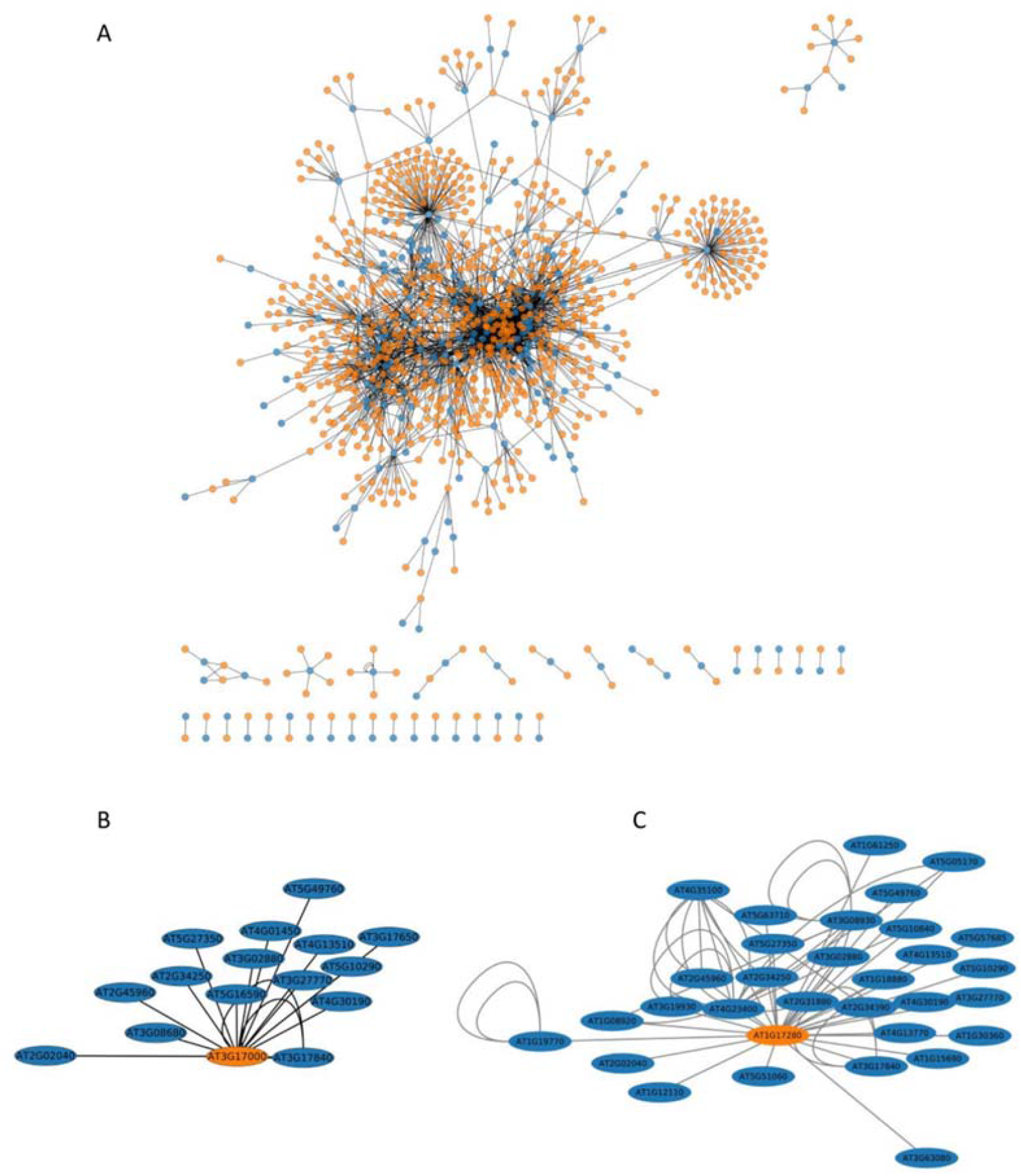
The interaction network of ubiquitinated proteins. lnteractants from a Y2H approach (Braun et al., 2011) and Split-Ub approach (Chen et al., 2012; Jones et al., 2014) were considered, and the network was visualized by Cytoscape (version 3.7.2). A. The network includes ubiquitinated proteins (blue) together with their reported interactants (orange) (Table 57a). B. The UBC32 subnetwork (see Table 57b). C. The UBC34 subnetwork (see Table S7c).

We then used a highly similar approach that focused specifically on DAUPs and their interactants, which made it possible to build another network of 326 proteins enriched in transporters and components of endocytic trafficking (Table S7d, Figure S7-S8). UBC32 and UBC34 appeared again as E2s targeting 9 proteins whose ubiquitination changed with mannitol treatment (Figure S7, Table 1). These proteins include NRT1;1-PTR8.3, NRT1-PTR6.3, two LRR-Receptor like kinases, a nodulin MtN21 like transporter, AHA2, the secretory carrier-associated membrane protein SCAMP3, the peroxidase GPX5, and an unknown protein.

### The osmotic phenotype of *ubc32, ubc33* and *ubc34* mutants

Root responses to osmotic stress involve high plasticity in root growth and architecture, which is partly determined by primary root (PR) growth. To reinforce the roles of UBC32 and UBC34 in the adaptive root response to osmotic stress, we studied the PR growth phenotype of corresponding mutants in control conditions and upon osmotic treatment. Since *AtUBC32* and *AtUBC34* belong to a small gene subfamily including *UBC33* (Ahn et al., 2018), we also studied the *ubc33* mutant and a triple knockout mutant line, *ubc32xubc33xubc34*, due to the putative redundancy between the 3 genes. Five-day-old plants were transferred to either control MS medium or MS medium supplemented with 0.2 M mannitol, and PR length was monitored for up to 5 days after transfer (Figure 6, Figure S9, Table S8). The root growth in control WT plants (Col) was inhibited by 14% one day after transfer, and by up to 43%, 5 days later (Figure 6). The *ubc32* and *ubc34* single mutants and the triple mutant showed a significantly higher PR growth inhibition ranging from 25 % at day 1 after transfer up to 50 % at day 5 after transfer (Figure 6). Thus, suppression of *AtUBC32* and *AtUBC34* favored PR growth inhibition under mannitol treatment, suggesting that these genes contribute to PR root growth under osmotic stress. This result is observable after only one day of treatment, suggesting that these genes play an early role in the response to osmotic stress.

**Figure 6.**
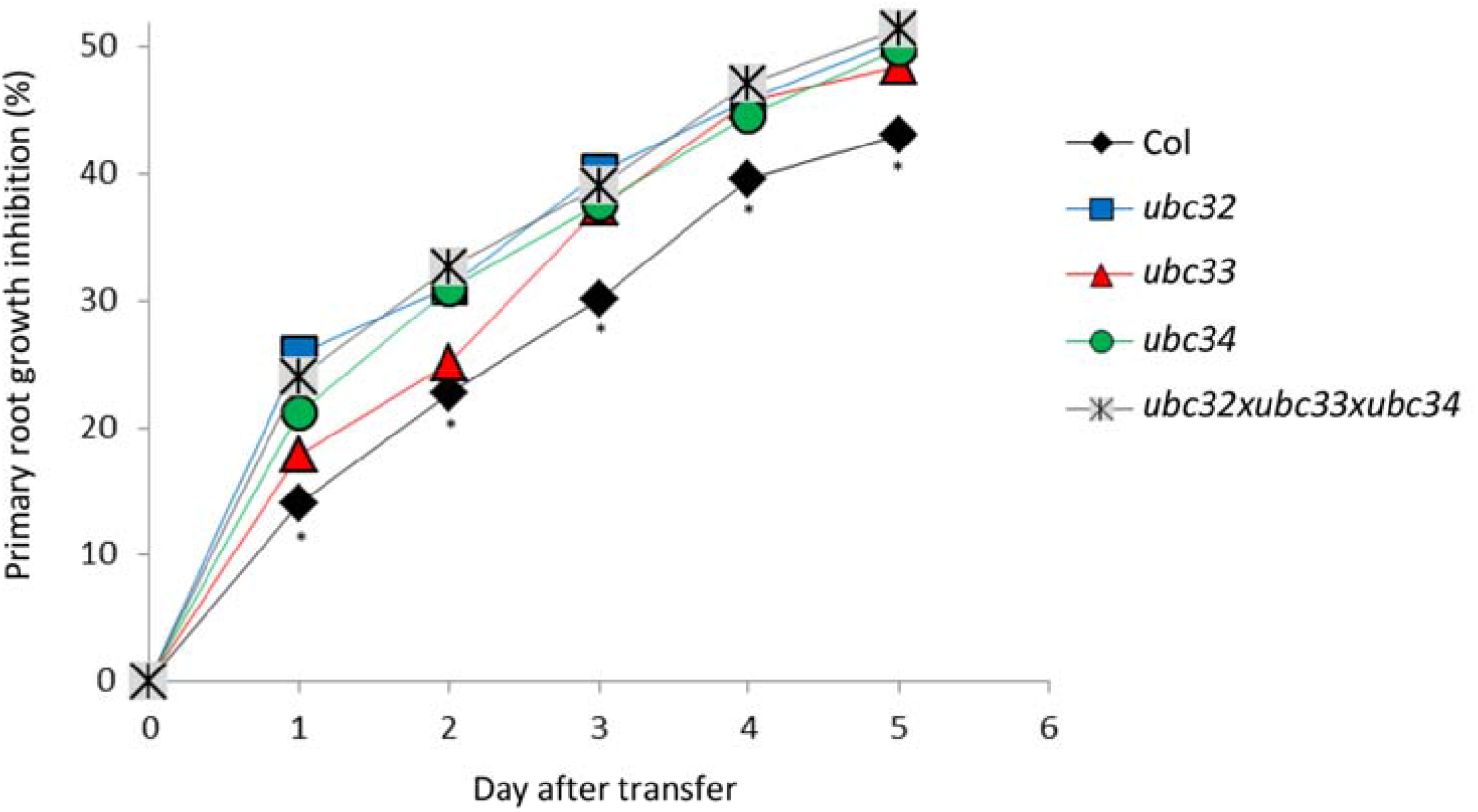
Inhibition of primary root length by mannitol in *Arabidopsis thaliana* WT plants (Col) and *ubc32, ubc33, ubc34* and *ubc32×ub33×ubc34* mutants. 15 plants per condition were grown for 5 days In MS medium and then transferred to MS medium and MS medium supplemented with 200 mM mannitol. Primary root length was monitored up to S days after transfer (Figure S9). Asterisks means that the VI/T Is significantly different from at least one mutant in a one-way ANOVA test combined with a Tukey test (p-value between 0.01 and 0.001 (Table S8)).

## Discussion

### A resource of ubiquitinated membrane proteins

Our study included 450 ubiquitinated proteins, significantly increasing the database size of ubiquitinated membrane protein substrates in plants. In comparison to other large-scale *Arabidopsis* ubiquitinomes (Maor et al., 2007; Manzano et al., 2008; Kim et al., 2013; Svozil et al., 2014; Walton et al., 2016; Zhang et al., 2019; Romero-Barrios et al., 2020; Grubb et al., 2021), 207 novel proteins, 90% of which are transmembrane proteins, were identified as ubiquitinated (Table S9). We also observed that 20% of Ubi-peptides can be additionally modified by oxidation, phosphorylation and acetylation (Table 1, Table S4), adding complexity to cellular signaling. The major proton pump AHA2 harbors 21 K-Ub sites (Tables S3), four of them showing quantitative variations upon mannitol treatment (Table 1). K888 in particular is ubiquitinated, and this site is located close to T881, whose phosphorylation leads to pump activation (Fuglsang et al., 2014). Moreover, the doubly modified peptide (*i*.*e*. by phosphorylation and ubiquitination) was accumulated upon mannitol treatment.

Recent studies have highlighted the importance of crosstalk between phosphorylation and ubiquitination in several plant signaling pathways (Vu et al., 2018), and the presence of such a K-Ub near a critical phospho-residue questions the role of this ubiquitination in the modulation of ATPase function.

Even though no ubiquitin motif could be described, the K-Ub residue appeared to be preferentially surrounded by an acidic residue (Figure 3). Similar observations can be obtained from ubiquitinome studies in petunia flower and rice embryo (Guo et al., 2017; He et al., 2020). By contrast, previous studies in rice leaf, wheat seedling and maize leaf have reported that alanine is enriched around K-Ub (Xie et al., 2015; Zhang et al., 2017; Wang et al., 2019). Thus, although no Ub motif could be identified, the presence of an acidic residue appears to be a common feature between 3 different tissues (root, flower, embryo), whereas the close presence of an alanine residue appears to be more specific for ubiquitinated residues in leaf proteins.

Our ubiquitinome study revealed that one-third of ubiquitinated proteins (151 proteins) harbor more than one ubiquitination site. In two recent studies, a total of 422 proteins were identified as carrying a K63-Ub linkage (Johnson and Vert, 2016; Romero-Barrios et al., 2020). Seventy-four of these proteins were identified in the present study, 29 of which were identified as carrying a unique K-Ub site that can be preferentially accounted for a K63-Ub linkage (Table S10). However, 44 proteins were also described in the present study as harboring at least 2 K-Ub sites (Table S10). This suggests that either all K-Ub residues are involved in K63-Ub linkage, or that different types of Ub linkages coexist within the same protein, including K63-Ub. This type of coexistence has already been shown in several studies; notably, the traffic and degradation of a particular receptor tyrosine kinase is likely to be regulated by different K-48 and K-63 poly-Ub editing mechanisms (Marx et al., 2010). Furthermore, the human nuclear protein Twist regulates a variety of cellular functions controlled by gene transcription events and can be ubiquitinated through K48-Ub linkage, which induces proteasome-dependent proteolysis and K63-Ub linkage for localization to the nucleus (Lee et al., 2016). Finally, in plants, the ubiquitination pattern of oleosin includes K48-Ub linkage that induces the proteasomal protein degradation, as well K63-Ub linkage which role remains unknown (Deruyffelaere et al., 2015). Thus, the specific fate of proteins can be dictated by specific coexisting poly-Ub linkages.

In addition to transporters and channels, we reported that vesicle transport-related proteins including clathrin assembly proteins, the AP-2 complex subunit, v-SNARE proteins, vesicle associated proteins and syntaxins were overrepresented in the ubiquitinome (Table S3). Endocytosis and endosomal trafficking are essential processes in cells for controlling the dynamics and turnover of plasma membrane proteins (Valencia et al., 2016). The recruitment of cargo into endocytic vesicles (e.g. clathrin-coated pits) involves the ‘endosomal sorting complex required for transport’ (ESCRT) multi-subunit complex, and requires adaptor proteins to eventually associate with clathrin (Valencia et al., 2016). The vesicles fuse with the acceptor organelle in a process mediated by factors such as SNAREs and small GTPases (Valencia et al., 2016). Here, we unexpectedly observed a concomitant ubiquitination of cargos and proteins from the endocytic machinery (Table S3). While a role for the ubiquitination of cargos in their endocytosis has recently emerged in plants (Romero-Barrios and Vert, 2018), the ubiquitination of proteins involved in endocytosis is rarely documented. In animals, several proteins involved in EGFR endocytosis were shown to be ubiquitinated after EGF stimulation (Haglund et al., 2002). In yeast, ubiquitination was recently shown to function as a recycling signal for sorting a SNARE into COPI vesicles in a non-degradative pathway (Xu et al., 2017). Therefore, our results suggest a role for ubiquitination in regulating the function of proteins involved in endocytic trafficking, highlighting a broad role for ubiquitin in internalizing and sorting cargo proteins.

### The role of ubiquitination in response to short-term osmotic treatment

Because ubiquitination can induce protein degradation, we looked for an inverse relationship between the abundance of proteins and their ubiquitinated form that could indicate a role for ubiquitination in protein degradation. Caution must be taken with this assumption, since it is not until the ubiquitin chain is assembled that it may act as a degradation signal (Clague et al., 2015). If a large proportion of ubiquitin is likely to be attached as mono-ubiquitin, this might skew the inverse relationship between the abundance of a protein and its ubiquitinated form. Although we may have overestimated the number of concerned proteins, only 10% of them (n=6) exhibited this inverse relationship, whereas 90% showed ubiquitination changes without any change in protein abundance (Table 1). Thus, for a majority of these membrane proteins, ubiquitination is not involved in protein degradation in response to short-term osmotic treatment. In particular, PIP2;1 abundance was unchanged upon short-term mannitol treatment, while its ubiquitination increased at K3 and K276 (Table 1). This osmotic treatment was shown to induce maximal *L*p_r_ inhibition by 60%, which can be accounted for by a decrease in aquaporin function (Di Pietro et al., 2013) and not PIP degradation, since the abundance of all PIPs was unchanged (Table S2). Thus, even though ubiquitination at K3 and K276 can be involved in PIP2;1 degradation (Figure 5, (Chen et al., 2021)), we assume that ubiquitination induces different consequences upon short-term mannitol treatment. Indeed, short-term osmotic treatment induces PIP2;1 selective endocytosis (Martiniere et al., 2019). In addition, PIP2;1 is ubiquitinated by K63-Ub linkage (Johnson and Vert, 2016), a poly-Ub linkage that plays a general function in the sorting of endocytosed cargos by the endosomal sorting complex required for transport (Romero-Barrios and Vert, 2018). Thus, we hypothesize that the increase in PIP2;1 ubiquitination induced by short-term treatment should participate in internalizing PIP2;1 and not in degradation of the protein. This result contrasts with the PIP2;1 degradation observed under long-term drought treatment, due to the simultaneous activity of UBC32 and the E3 ligase Rma1 in ubiquitinating PIP2;1 at K276 (Chen et al., 2021). The pairing of E3s with different E2s is dynamic and changes in response to external stimuli (Turek et al., 2018). Thus, under short-term osmotic stress, a specific E2/E3 combination that differs from UBC32/Rma1 could regulate PIP2;1 internalization. However, this hypothesis will require additional experiments.

### UBC32 and UBC34 contribute to primary root growth under osmotic stress

E2s have recently emerged as key mediators of chain assembly, in particular by dictating the K residue within ubiquitin used to link the moieties in a chain (Turek et al., 2018). Our protein-protein network analysis identified UBC32 and UBC34 as E2s that putatively interact with proteins whose ubiquitination changed upon short mannitol treatment (Table 1, Figure S7). Recent expression studies have suggested that E2s participate in abiotic stress responses (Zhou et al., 2010; Zhiguo et al., 2015; Sharma and Bhatt, 2018). Surprisingly, the role of *UBC32* appears contradictory, since it has been described as playing both negative and positive roles in response to long-term drought in (Ahn et al., 2018) and (Chen et al., 2021), respectively. In particular, the survival rate of 3-week-old *ubc32* plants after 12-20 days of withholding water has been reported to be both higher and lower than WT plants in (Ahn et al., 2018) and (Chen et al., 2021), respectively. Our study reveals that, in the context of short-term osmotic treatment, UBC32 and UBC34 positively regulate PR growth, thus playing a positive role in osmotic stress tolerance. *UBC32, UBC33* and *UBC34* are all reported to participate in the endoplasmic reticulum-associated degradation (ERAD) pathway in *Arabidopsis*, which is a major degradation system involved in removing misfolded or unfolded proteins retained in the ER (Cui et al., 2012; Chen et al., 2016; Chen et al., 2017; Zheng et al., 2019; Chen et al., 2021; Zhang et al., 2021). The involvement of ERAD components suggests that short-term osmotic stress may also result in ER/protein stress, which engages ERAD to control the secretion of plasma membrane proteins. However, upon 1-h mannitol treatment, proteins that putatively interact with UBC32 and UBC34 and show increased ubiquitination did not display any decreased cellular abundance (Table 1). We thus speculate that UBC32 and UBC34 are not simply ERAD components, but that they also participate in the ubiquitination process in other subcellular organelles such as the plasma membrane in response to a short-term osmotic stress. In particular, an internalization of the aquaporin PIP2;1 (Chen et al., 2021) was observed upon a short-term osmotic stress (Martiniere et al., 2019). We thus speculate that, upon a short-term osmotic stress, a rapid coordinated internalization of aquaporins and of ions transporters involved in plant mineral nutrition could contribute to maintain a minimal root growth. Conversely, absence of internalization would impair root development. Soil is extremely heterogeneous and root growth maintenance under unfavorable local environment could allow the root tip to reach a more favorable environment and then to maintain on a longer term root foraging and plant survival.

## Conclusion

Our data present the ubiquitinome of root membrane proteins and its variation under osmotic stress. Importantly, the results highlight specific post-translational modification patterns and suggest approaches for exploring the physiological role of lysine ubiquitination in plants under osmotic stress. Our results open new perspectives in the involvement of ubiquitination and trafficking of root plasma membrane transporters in response to changes in local environment and future studies exploring the function of these ubiquitination in PIP2;1 and ion transporters will be necessary to delineate its role in the root nutrient foraging.

## Material and methods

### Plant materials and growth conditions

*Arabidopsis thaliana* ecotype Columbia (Col-8) was used as the control wild type (WT) plant. The *atubc32* (SALK_092817), *atubc33 (GABI_105_D10), atubc34* (SAIL_1249_C08) single T-DNA insertion mutant alleles and the triple mutant *atubc32*x*atubc33*x*atubc34* were obtained from (Ahn et al., 2018). Homozygosity of these mutants was verified using primers described in (Ahn et al., 2018). For proteome and ubiquitinome studies, WT seeds were surface-sterilized and sown in 0.2-mL tubes containing 0.8% agar prepared in pH 5.5 Hoagland-based solution (0.5 mM KH_2_PO_4_, 1.25 mM KNO_3_, 0.75 mM MgSO_4_, 1.5 mM CaNO_3_, 50 μM H_3_BO_3_, 0.7 μM CuSO_4_, 1 μM ZnSO_4_, 12 μM MnSO_4_, 0.24 μM Na2MoO_4_, 50 μM Fe^3+^-EDTA). After 7 days in the growth chamber, the bottoms of the tubes were cut off prior to transfer in 2.5-L opaque recipients in the same medium. Plants were grown for 8 weeks under short-day conditions (8 h/16 h day/night; 23°C/20°C day/night) at a light intensity of 160 μmol.m^-2^.s^-1^ and 65% humidity. Plants were treated with 0.2 M mannitol for 1 h. Roots were harvested and stored at -80°C until analysis.

### Microsome extraction

Roots were crushed with a PULVERISETTE 2 Mortar Grinder (Fritsch) in liquid nitrogen and microsomal proteins were extracted according to (Di Pietro et al., 2013), except that the grinding buffer contained 10 mM N-ethylmaleimide and that the pellets were resuspended with a potter in 200 μL of Laemmli 1X buffer (65 mM Tris–HCl, pH 7.5, 5% glycerol, 2% SDS) (Laemmli, 1970). Proteins were quantified using a detergent compatible with the Bradford assay kit (Thermo Scientific).

### Protein digestion

For proteome analysis, three independent biological replicates from the control condition and mannitol-treated plants were used. 10 μg of microsomes were fractionated using 10% precast SDS-PAGE gel electrophoresis (Bio-Rad). After staining with Coomassie blue (R250, Bio-Rad), the gel was rinsed with acetic acid/methanol (Destain, Bio-Rad). Each lane was cut into 4 bands. For the ubiquitinome study, 2 independent biological replicates from the control condition and mannitol-treated plants were used, and microsomal fractions (1.2 mg) were subjected to an in-tube acrylamide inclusion (13% acrylamide/bis-acrylamide, 0.6% ammonium persulfate, 2.5% TEMED) adapted from (Balliau et al., 2018). For proteome and ubiquitinome analyses, gel slices were treated according to (Chen et al., 2019), with the exception that proteins were alkylated with 50 mM chloroacetamide for ubiquitinome experiments. Proteins were digested with trypsin (Sequencing Grade Modified Trypsin, Promega, Madison, WI, USA) at a 1:50 (trypsin:protein) ratio at 37°C overnight. Peptides were extracted according to (Chen et al., 2019).

### Enrichment of ubiquitinated peptides

Tryptic peptides were filtered through a C18 cartridge (Sep-Pack Classic, Waters) equilibrated with 0.1% TFA. After loading on the column, the peptides were washed twice with 0.1% TFA and then with 0.1% TFA and 5% ACN. Peptides were eluted with 0.1% TFA and 40% ACN, pooled, frozen overnight at -80°C, and finally evaporated. Immunoprecipitation experiments were performed with 15 μL of Pan anti-glycine lysine antibody conjugated to agarose beads (PTM Biolabs, Chicago, IL, USA) according to the manufacturer’s instructions. Briefly, tryptic peptides were dissolved in 300 μL of WASH I (100 mM NaCl, 1 mM EDTA, 20 mM Tris-HCl, 0.25% n-Dodecyl β-D-maltoside, pH 8.0), incubated 4 h at room temperature on a rotary shaker, and then sequentially washed 3 times with WASH I, 3 times with WASH II (100 mM NaCl, 1 mM EDTA, 20 mM Tris-HCl, pH 8.0), and 3 times with HPLC-grade water. The elution was performed 3 times with 100 μL of 0.1% TFA. For the LC-MS/MS experiment, 300 μL of pooled eluates were dried under vacuum centrifuge and resuspended in 2% FA.

### LC-MS/MS analysis

The LC-MS/MS experiments were performed using a NCS 3500RS-ProFlow nano system (Thermo Fisher Scientific Inc., Waltham, MA, USA) interfaced online with a nano easy ion source and a Q-Exactive Plus Orbitrap mass spectrometer (Thermo Fisher Scientific Inc, Waltham, MA, USA). The samples were analyzed in a data-dependent acquisition mode. For total proteome and ubiquitinome experiments, 2 μL and 6 μL of peptides were injected, respectively. Peptides were first loaded onto a pre-column (Thermo Scientific PepMap 100 C18, 5 μm particle size, 100 Å pore size, 300 μm i.d. x 5 mm length) from the Ultimate 3000 autosampler with 0.05% TFA in water at a flow rate of 10 μL/min. The peptides were separated by reverse-phase column (Thermo Scientific PepMap C18, 3 μm particle size, 100 Å pore size, 75 μm i.d. x 50 cm length) at a flow rate of 300 nL/min. After a 3-min loading period, the column valve was switched to allow elution of peptides from the pre-column onto the analytical column. The loading buffer (solvent A) consisted of 0.1% FA in water, and the elution buffer (solvent B) was 0.1% FA in 80% ACN. The employed 3-step gradient consisted of 4-25% of solvent B until 50 min for ubiquitinome (103 min for total proteome), then 25-40% of solvent B from 50 to 60 min for ubiquitinome (from 103 to 123 min for total proteome), and finally 40-90% of solvent B from 60 to 62 min (123 to 125 min for total proteome). The total run time was 90 min for ubiquitinome (150 min for total proteome), including a high organic wash step and a re-equilibration step. Peptides were transferred to the gaseous phase with positive ion electrospray ionization at 1.8 kV. In the data-dependent acquisition procedure, the top 10 precursors were acquired between 375 and 1,500 m/z with a 2 Th (Thomson) selection window, a dynamic exclusion of 40 s, a normalized collision energy of 27, and resolutions of 70,000 for MS and 17,500 for MS2. Spectra were recorded with Xcalibur software (4.3.31.9) (Thermo Fisher Scientific). The mass spectrometry proteomics data were deposited at the ProteomeXchange Consortium *via* the PRIDE partner repository with the dataset identifier PXD022249. The reviewer account details are: Username, Password: B1h58X9Q.

### Identification and quantification of whole proteome and ubiquitinome

For the proteome and the ubiquitinome, the resulting MS/MS data were processed using MaxQuant with an integrated Andromeda search engine (version 1.6.6.0). Tandem mass spectra were searched against the TAIR10 database (35,417 entries). The minimal peptide length was set to 6. The criteria “Trypsin/P” was chosen as the digestion enzyme. Carbamidomethylation of cysteine was selected as fixed modification and methionine oxidation, N-terminal acetylation and phosphorylation (S/T/Y) were systematically selected as variable modifications. Up to 4 missed cleavages were systematically allowed.

For proteome analysis, the mass tolerance of the precursor was 20 and 4.5 ppm for the first and main searches, respectively, and was 20 ppm for the fragment ions. The function “match between run” was used. Proteins were identified provided that they contained one unique trypsin peptide. The rates of false peptide sequence assignment and false protein identification were fixed to be lower than 1%. Quantification was performed with at least 2 peptides per protein, one of them unique to the protein. To investigate differentially expressed proteins, Student’s t-test was performed using protein Label-Free Quantification (LFQ) intensity values when present in at least 2 replicates and in at least 2 biological replicates per condition.

For ubiquitinome analysis, “GlyGly” on K residue was additionally specified as a variable modification. The function “match between run” was not applied. The minimum score for peptides was set to 40. The intensity of each peptide from the “evidence” table was normalized to the sum of all peptide intensities in each sample, and a t-test was performed to investigate differentially-expressed peptides. The ubiquitinated peptides with consistent fold changes in two replicates were counted, and the significance of the abundance change among samples was evaluated as differentially expressed by a Student’s t-test. A p-value < 0.05 was considered statistically significant. The appearance/disappearance of peptides was considered on condition of their presence in two biological replicates and the corresponding absence from the two other biological replicates. We defined “absence” as no razor or unique peptide in any biological condition replicate, and “presence” as the identification of at least one unique peptide in all replicates of a biological condition.

### Bioinformatic analyses

Gene Ontology (GO) term association and enrichment analyses were performed using Panther (http://www.pantherdb.org/) (Mi et al., 2013). Fold enrichments were calculated based on the frequency of proteins annotated to the term compared with their frequency in the *Arabidopsis* proteome. The p-value combined with the false discovery rate (FDR) correction was used as criteria of significant enrichment for GO catalogs, whereas a p-value < 0.05 was considered to be enriched for GO terms. The most specific subclasses were considered. The GO annotation was classified based on the “biological processes”, “molecular functions” and “cellular components” categories. GO terms were reduced with rrvgo (https://ssayols.github.io/rrvgo/). The number of transmembrane domains was estimated with Aramemnon (http://aramemnon.botanik.uni-koeln.de/). The p-logo software (O’Shea et al., 2013) (https://plogo.uconn.edu/) was used to analyze the models of the sequences with amino acids in specific positions of ubiquitin-21-mers (10 amino acids upstream and downstream of the K-Ub site) in all of the protein sequences. In addition, the *Arabidopsis* proteome was used as the background database, and the other parameters were set to default values. Protein-protein interaction data were obtained from plant interactome databases, including results from a yeast two-hybrid approach (Braun et al., 2011) and from Split-ubiquitin approaches (Chen et al., 2012; Jones et al., 2014), in order to build a network including these ubiquitinated proteins together with their reported interactants. Protein-protein interaction networks were visualized using Cytoscape version 3.7.2 (Shannon et al., 2003).

### Ectopic expression of PIP2;1 in suspension cells

Mutated PIP2;1 cDNAs were constructed according to (Santoni et al., 2006). Biolistic transformation of 5-day-old suspension cells was performed as described in (Santoni et al., 2006), and transformed cells were selected on 50 mg/L of hygromycin. Briefly, independent transformed cells were isolated and further cultured on 40 mg/L of hygromycin. Stable insertion of the T-DNA was checked by PCR, and the expression of PIP was detected by western blot as described in (Santoni et al., 2006). Extraction of total proteins from suspension cells and ELISA measurements of PIP2 abundance were performed as described in (Santoni et al., 2006).

### Root architecture analyses

Plants were stratified for 2 days at 4°C and grown vertically on agar plates containing half-strength Murashige and Skoog (MS) medium supplemented with 1% (w/v) sucrose and 2.5 mM MES-KOH pH 6, in a self-contained imaging unit equipped with a 16M pixel linear camera, a telecentric objective and collimated LED backlight. Plants were grown in the imaging automat dedicated growth chamber at 23°C in a 16-h light/8-h dark cycle with 70% relative humidity and a light intensity of 185 μmol.m^-2^.s^-1^ (Vegeled Floodlight, Colasse Seraing, Belgium). Plates were imaged every day for 5 days.

## Abbreviations

K-Ub: ubiquitinated lysine
LC-MS/MS: liquid chromatography-tandem mass spectrometry
*L*p_r_: root hydraulic conductivity
PIP: plasma membrane intrinsic protein
Ub: ubiquitin
Ubi-peptides: ubiquitinated peptides

## Supplemental material

Supplemental Figure S1. Characterization of the microsomal fraction.

Supplemental Figure S2. Functional enrichment analysis of differentially accumulated proteins (DAPs) in response to mannitol.

Supplemental Figure S3. Types of Ub linkages.

Supplemental Figure S4. Functional enrichment analysis of differentially accumulated ubiquitinated proteins (DAUPs) in response to mannitol.

Supplemental Figure S5: K-Ub residues in PIP aquaporins.

Supplemental Figure S6: Functional enrichment analysis of the interactome of ubiquitinated proteins (corresponding to Figure 6).

Supplemental Figure S7: Interaction network of DAUPs.

Supplemental Figure S8: Functional enrichment analysis of the DAUP interactome.

Supplemental Figure S9: Root growth phenotype of WT plants and *ubc* mutants in 3-day control and 0.2 M mannitol conditions.

Supplemental Table S1: The root microsomal proteome of plants in the control condition and upon mannitol treatment.

Supplemental Table S2: Proteins with quantitative variations upon mannitol treatment.

Supplemental Table S3: Inventory of ubiquitinated proteins in a root microsomal fraction.

Supplemental Table S4: Inventory of Ubi-peptides in root microsomes.

Supplemental Table S5: Determination of the ubiquitination motif.

Supplemental Table S6: Peptide quantification after immunopurification with the anti-KGG antibody.

Supplemental Table S7a: Interaction network of ubiquitinated proteins.

Supplemental Table S7b: Interaction network of UBC32.

Supplemental Table S7c: Interaction network of UBC34.

Supplemental Table S7d: Interaction network of DAUPs.

Supplemental Table S8a: One-way analysis of variance of root growth inhibition upon 1-day treatment with 0.2 M mannitol (corresponding to Figure 6).

Supplemental Table S8b: One-way analysis of variance of root growth inhibition upon 2-day treatment with 0.2 M mannitol (corresponding to Figure 6).

Supplemental Table S8c: One-way analysis of variance of root growth inhibition upon 3-day treatment with 0.2 M mannitol (corresponding to Figure 6).

Supplemental Table S8d: One-way analysis of variance of root growth inhibition upon 4-day treatment with 0.2 M mannitol (corresponding to Figure 6).

Supplemental Table S8e: One-way analysis of variance of root growth inhibition upon 5-day treatment with 0.2 M mannitol (corresponding to Figure 6).

Supplemental Table S9a: Ubiquitinated proteins described in other Arabidopsis ubiquitome studies. Table S9b: Novel ubiquitinated proteins identified in the present work.

Supplemental Table S10: Ubiquitinated proteins identified in the present work and described in the literature as containing a K63-Ub linkage.

## Acknowledgments

We thank Dr. Ahn (Yonsei University, Seoul, Republic of Korea) for providing seeds of mutant *ubc32, ubc33, ubc34* plants and the triple mutant *ubc32xubc33xubc34*. We thank Brandon Loveall of Improvence for English proofreading of the manuscript.

*Conflict of interest statement*. The authors declare no conflicts of interest.

## Data availability statement

The mass spectrometry proteomics data were deposited at the ProteomeXchange Consortium via the PRIDE partner repository with the dataset identifier PXD022249.

## Parsed Citations

Ahn MY, Oh TR, Seo DH, Kim JH, Cho NH, Kim WT (2018) Arabidopsis group XIV ubiquitin-conjugating enzymes AtUBC32, AtUBC33, and AtUBC34 play negative roles in drought stress response. Journal of Plant Physiology 230: 73–79

Balliau T, Blein-Nicolas M, Zivy M (2018) Evaluation of Optimized Tube-Gel Methods of Sample Preparation for Large-Scale Plant Proteomics. Proteomes 6

Braun P, Carvunis AR, Charloteaux B, Dreze M, Ecker JR, Hill DE, Roth FP, Vidal M, Galli M, Balumuri P, Bautista V, Chesnut JD, Kim RC, de los Reyes C, Gilles P, Kim CJ, Matrubutham U, Mirchandani J, Olivares E, Patnaik S, Quan R, Ramaswamy G, Shinn P, Swamilingiah GM, Wu S, Byrdsong D, Dricot A, Duarte M, Gebreab F, Gutierrez BJ, MacWilliams A, Monachello D, Mukhtar MS, Poulin MM, Reichert P, Romero V, Tam S, Waaijers S, Weiner EM, Cusick ME, Tasan M, Yazaki J, Ahn YY, Barabasi AL, Chen HM, Dangl JL, Fan CY, Gai LT, Ghoshal G, Hao T, Lurin C, Milenkovic T, Moore J, Pevzner SJ, Przulj N, Rabello S, Rietman EA, Rolland T, Santhanam B, Schmitz RJ, Spooner W, Stein J, Vandenhaute J, Ware D, Arabidopsis Interactome Mapping C (2011) Evidence for network evolution in an Arabidopsis Interactome map. Science 333: 601–607

Callis J (2014) The Ubiquitination Machinery of the Ubiquitin System Arabidopsis book 12

Chen J, Lalonde S, Obrdlik P, Vatani AN, Parsa SA, Vilarino C, Revuelta JL, Frommer WB, Rhee SY (2012) Uncovering Arabidopsis membrane protein interactome enriched in transporters using mating-based split ubiquitin assays and classification models. Frontiers Plant Sci. doi: 10.3389/fpls.2012.00124

Chen Q, Liu R, Wu Y, Wei S, Wang Q, Zheng Y, Ran X, Shang X, Yu F, Yang X, Liu L, Huang X, Wang Y, Xie Q (2021) ERAD-related E2 and E3 enzymes modulate the drought response by regulating the stability of PIP2 aquaporins. The Plant Cell 33: 2883–2898

Chen Q, Liu RJ, Wang Q, Xie Q (2017) ERAD Tuning of the HRD1 Complex Component AtOS9 Is Modulated by an ER-Bound E2, UBC32. Molecular Plant 10: 891–894

Chen Q, Zhong YW, Wu YR, Liu LJ, Wang PF, Liu RJ, Cui F, Li QL, Yang XY, Fang SY, Xie Q (2016) HRD1-mediated ERAD tuning of ER-bound E2 is conserved between plants and mammals. Nature Plants 2

Chen Y, Rofidal V, Hem S, Gil J, Nosarzewska J, Berger N, Demolombe V, Bouzayen M, Azhar BJ, Shakeel SN, Schaller GE, Binders BM, Santoni V, Chervin C (2019) Targeted Proteomics Allows Quantification of Ethylene Receptors and Reveals SIETR3 Accumulation in Never-Ripe Tomatoes. Frontiers in Plant Science 10

Clague MJ, Heride C, Urbe S (2015) The demographics of the ubiquitin system. Trends in Cell Biology 25: 417–426

Cui F, Liu LJ, Li QL, Yang CW, Xie Q (2012) UBC32 Mediated Oxidative Tolerance in Arabidopsis. Journal of Genetics and Genomics 39: 415–417

Deruyffelaere C, Bouchez I, Morin H, Guillot A, Miquel M, Froissard M, Chardot T, D’Andrea S (2015) Ubiquitin-Mediated Proteasomal Degradation of Oleosins is Involved in Oil Body Mobilization During Post-Germinative Seedling Growth in Arabidopsis. Plant and Cell Physiology 56: 1374–1387

Di Pietro M, Vialaret J, Li G-W, Hem S, Prado K, Rossignol M, Maurel C, Santoni V (2013) Coordinated post-translational responses of aquaporins to abiotic and nutritional stimuli in Arabidopsis roots. Molecular and Cellular Proteomics 12: 3886–3897

Falhof J, Pedersen JT, Fuglsang AT, Palmgren M (2016) Plasma Membrane H+-ATPase Regulation in the Center of Plant Physiology. Molecular Plant 9: 323–337

Fuglsang AT, Kristensen A, Cuin TA, Schulze WX, Persson J, Thuesen KH, Ytting CK, Oehlenschlaeger CB, Mahmood K, Sondergaard TE, Shabala S, Palmgren MG (2014) Receptor kinase-mediated control of primary active proton pumping at the plasma membrane. Plant Journal 80: 951–964

Grubb LE, Derbyshire P, Dunning KE, Zipfel C, Menke FLH, Monaghan J (2021) Large-scale identification of ubiquitination sites on membrane-associated proteins in Arabidopsis thaliana seedlings. Plant Physiology 185: 1483–1488

Guo JH, Liu JX, Wei Q, Wang RM, Yang WY, Ma YY, Chen GJ, Yu YX (2017) Proteomes and Ubiquitylomes Analysis Reveals the Involvement of Ubiquitination in Protein Degradation in Petunias. Plant Physiology 173: 668–687

Guo QF, Zhang J, Gao Q, Xing SC, Li F, Wang W (2008) Drought tolerance through overexpression of monoubiquitin in transgenic tobacco. Journal of Plant Physiology 165: 1745–1755

Haglund K, Shimokawa N, Szymkiewicz I, Dikic I (2002) Cbl-directed monoubiquitination of CIN85 is involved in regulation of ligand-induced degradation of EGF receptors. Proceedings of the National Academy of Sciences of the United States of America 99: 12191–12196

He DL, Li M, Damaris RN, Bu C, Xue JY, Yang PF (2020) Quantitative ubiquitylomics approach for characterizing the dynamic change and extensive modulation of ubiquitylation in rice seed germination. Plant Journal 101: 1430–1447

Johanson U, Gustavsson S (2002) Anew subfamily of major intrinsic proteins in plants. Molecular Biology and Evolution 19: 456–461

Johnson A, Vert G (2016) Unraveling K63 Polyubiquitination Networks by Sensor-Based Proteomics. Plant Physiology 171: 1808–1820

Jones AM, Xuan YH, Xu M, Wang RS, Ho CH, Lalonde S, You CH, Sardi MI, Parsa SA, Smith-Valle E, Su TY, Frazer KA, Pilot G, Pratelli R, Grossmann G, Acharya BR, Hu HC, Engineer C, Villiers F, Ju CL, Takeda K, Su Z, Dong QF, Assmann SM, Chen J, Kwak JM, Schroeder JI, Albert R, Rhee SY, Frommer WB (2014) Border control - A membrane-linked interactome of Arabidopsis. Science 344: 711–716

Kang HH, Zhang M, Zhou SM, Guo QF, Chen FJ, Wu JJ, Wang W (2016) Overexpression of wheat ubiquitin gene, Ta-Ub2, improves abiotic stress tolerance of Brachypodium distachyon. Plant Science 248: 102–115

Kim DY, Scalf M, Smith LM, Vierstra RD (2013) Advanced proteomic analyses yield a deep catalog of ubiquitylation targets in Arabidopsis. Plant Cell 25: 1523–1540

Komander D, Rape M (2012) The Ubiquitin Code. In RD Kornberg, ed, Annual Review of Biochemistry, Vol 81, Vol 81. Annual Reviews, Palo Alto, pp 203–229

Laemmli UK (1970) Cleavage of structural proteins during the assembly of the head of bacteriophage T4. Nature 222: 680–865

Lee HJ, Li CF, Ruan DN, Powers S, Thompson PA, Frohman MA, Chan CH (2016) The DNADamage Transducer RNF8 Facilitates Cancer Chemoresistance and Progression through Twist Activation. Molecular Cell 63: 1021–1033

Liu TY, Huang TK, Tseng CY, Lai YS, Lin SI, Lin WY, Chen JW, Chiou TJ (2012) PHO2-Dependent Degradation of PHO1 Modulates Phosphate Homeostasis in Arabidopsis. Plant Cell 24: 2168–2183

Lobell DB, Schlenker W, Costa-Roberts J (2011) Climate Trends and Global Crop Production Since 1980. Science 333: 616–620

Manzano C, Abraham Z, Lopez-Torrejon G, Del Pozo JC (2008) Identification of ubiquitinated proteins in Arabidopsis. Plant Molecular Biology 68: 145–158

Maor R, Jones A, Nuhse TS, Studholme DJ, Peck SC, Shirasu K (2007) Multidimensional protein identification technology (MudPIT) analysis of ubiquitinated proteins in plants. Molecular & Cellular Proteomics 6: 601–610

Martiniere A, Fiche JB, Smokvarska M, Mari S, Alcon C, Dumont X, Hematy K, Jaillais Y, Nollmann M, Maurel C (2019) Osmotic Stress Activates Two Reactive Oxygen Species Pathways with Distinct Effects on Protein Nanodomains and Diffusion. Plant Physiology 179: 1581–1593

Marx C, Held JM, Gibson BW, Benz CC (2010) ErbB2 Trafficking and Degradation Associated with K48 and K63 Polyubiquitination. Cancer Research 70: 3709–3717

Maurel C, Boursiac Y, Luu D-T, Santoni V, Shahzad Z, Verdoucq L (2015) Aquaporins in Plants. Physiological reviews 95: 1321–1358

Mi HY, Muruganujan A, Casagrande JT, Thomas PD (2013) Large-scale gene function analysis with the PANTHER classification system. Nature Protocols 8: 1551–1566

Mukhopadhyay D, Riezman H (2007) Proteasome-independent functions of ubiquitin in endocytosis and signaling. Science 315: 201–205

Nelson CJ, Millar AH (2015) Protein turnover in plant biology. Nature Plants 1

O’Shea JP, Chou MF, Quader SA, Ryan JK, Church GM, Schwartz D (2013) pLogo: a probabilistic approach to visualizing sequence motifs. Nature methods 10: 1211-+

Pan WB, Lin BY, Yang XY, Liu LJ, Xia R, Li JG, Wu YR, Xie Q (2020) The UBC27-AIRP3 ubiquitination complex modulates ABA signaling by promoting the degradation of ABI1 in Arabidopsis. Proceedings of the National Academy of Sciences of the United States of America 117: 27694–27702

Pickart CM, Eddins MJ (2004) Ubiquitin: structures, functions, mechanisms. Biochim. Biophys. Acta-Molecular Cell Research 1695: 55–72

Quigley F, Rosenberg JM, Shachar-Hill Y, Bohnert HJ (2002) From genome to function: the Arabidopsis aquaporins. Genome Biology 3: 1–17

Romero-Barrios N, Monachello D, Dolde U, Wong A, San Clemente H, Cayrel A, Johnson A, Lurin C, Vert G (2020) Advanced Cataloging of Lysine-63 Polyubiquitin Networks by Genomic, Interactome, and Sensor-Based Proteomic Analyses. Plant Cell 32: 123–138

Romero-Barrios N, Vert G (2018) Proteasome-independent functions of lysine-63 polyubiquitination in plants. New Phytologist 217: 995–1011

Santoni V, Verdoucq L, Sommerer N, Vinh J, Pflieger D, Maurel C (2006) Methylation of aquaporins in plant plasma membrane. Biochem. J. 400: 189–197

Shannon P, Markiel A, Ozier O, Baliga NS, Wang JT, Ramage D, Amin N, Schwikowski B, Ideker T (2003) Cytoscape: Asoftware environment for integrated models of biomolecular interaction networks. Genome Res. 13: 2498–2504

Sharma B, Bhatt TK (2018) Genome-wide identification and expression analysis of E2 ubiquitin-conjugating enzymes in tomato (vol 7, 8613, 2017). Scientific Reports 8

Stone SL (2018) Role of the Ubiquitin Proteasome System in Plant Response to Abiotic Stress. In L Galluzzi, ed, International Review of Cell and Molecular Biology, Vol 343, Vol 343. Academic Press Ltd-Elsevier Science Ltd, London, pp 65–110

Svozil J, Hirsch-Hoffmann M, Dudler R, Gruissem W, Baerenfaller K (2014) Protein Abundance Changes and Ubiquitylation Targets Identified after Inhibition of the Proteasome with Syringolin A. Molecular & Cellular Proteomics 13: 1523–1536

Tsuchiya H, Tanaka K, Saeki Y (2013) The parallel reaction monitoring method contributes to a highly sensitive polyubiquitin chain quantification. Biochem Biophys Res Commun 436: 223–229

Turek I, Tischer N, Lassig R, Trujillo M (2018) Multi-tiered pairing selectivity between E2 ubiquitin-conjugating enzymes and E3 ligases. Journal of Biological Chemistry 293: 16324–16336

Valencia JP, Goodman K, Otegui MS (2016) Endocytosis and Endosomal Trafficking in Plants. In SS Merchant, ed, Annual Review of Plant Biology, Vol 67, Vol 67. Annual Reviews, Palo Alto, pp 309–335

Vu LD, Gevaert K, De Smet I (2018) Protein Language: Post-Translational Modifications Talking to Each Other. Trends in Plant Science 23: 1068–1080

Walsh CK, Sadanandom A(2014) Ubiquitin chain topology in plant cell signaling: a new facet to an evergreen story. Front. Plant Sci. 5: 122

Walton A, Stes E, Cybulski N, Van Bel M, Inigo S, Durand AN, Timmerman E, Heyman J, Pauwels L, De Veylder L, Goossens A, De Smet I, Coppens F, Goormachtig S, Gevaert K (2016) It’s Time for Some “Site”-Seeing: Novel Tools to Monitor the Ubiquitin Landscape in Arabidopsis thaliana. Plant Cell 28: 6–16

Wang YF, Chao Q, Li Z, Lu T-C, Zheng H-Y, Zhao C-F, Shen Z, Li X-H, Wang B-C (2019) Large-scale Identification and Time-course Quantification of Ubiquitylation Events During Maize Seedling De-etiolation. Genomics Proteomics Bioinformatics 17: 603–622

Xie X, Kang HX, Liu WD, Wang GL (2015) Comprehensive Profiling of the Rice Ubiquitome Reveals the Significance of Lysine Ubiquitination in Young Leaves. Journal of Proteome Research 14: 2017–2025

Xu P, Hankins HM, MacDonald C, Erlinger SJ, Frazier MN, Diab NS, Piper RC, Jackson LP, MacGurn JA, Graham TR (2017) COPI mediates recycling of an exocytic SNARE by recognition of a ubiquitin sorting signal. Elife 6

Zhang L, Yu ZP, Xu Y, Yu M, Ren Y, Zhang SZ, Yang GD, Huang JG, Yan K, Zheng CC, Wu CG (2021) Regulation of the stability and ABA import activity of NRT1.2/NPF4.6 by CEPR2-mediated phosphorylation in Arabidopsis. Molecular Plant 14: 633–646

Zhang N, Xu J, Liu XY, Liang WX, Xin MM, D.JK, Hu ZR, Peng HR, Guo WL, Ni ZF, Sun QX, Yao YY (2019) Identification of HSP90C as a substrate of E3 ligase TaSAP5 through ubiquitylome profiling. Plant Science 287

Zhang N, Zhang LR, Shi CN, Tian QZ, Lv GG, Wang Y, Cui DQ, Chen F (2017) Comprehensive profiling of lysine ubiquitome reveals diverse functions of lysine ubiquitination in common wheat. Scientific Reports 7

Zheng L, Chen YJ, Ding D, Zhou Y, Ding LP, Wei JH, Wang HZ (2019) Endoplasmic reticulum-localized UBC34 interaction with lignin repressors MYB221 and MYB156 regulates the transactivity of the transcription factors in Populus tomentosa. BMC Plant Biology 19

Zhiguo E, Zhang YP, Li TT, Wang L, Zhao HM (2015) Characterization of the Ubiquitin-Conjugating Enzyme Gene Family in Rice and Evaluation of Expression Profiles under Abiotic Stresses and Hormone Treatments. Plos One 10

Zhou GA, Chang RZ, Qiu LJ (2010) Overexpression of soybean ubiquitin-conjugating enzyme gene GmUBC2 confers enhanced drought and salt tolerance through modulating abiotic stress-responsive gene expression in Arabidopsis. Plant Molecular Biology 72: 357–367

